# Parental age at conception on mouse lemur’s offspring longevity: sex-specific maternal effects

**DOI:** 10.1101/2022.03.09.483593

**Authors:** Perret Martine, Anzereay Aude

## Abstract

Parental age at conception often influences offspring’s longevity, a phenomenon referred as the “Lansing effect” described in large variety of organisms. But, the majority of the results refer to the survival of juveniles, mainly explained by an inadequate parental care by the elderly parents, mostly the mothers. Studies on the effect of parental age on offspring’s longevity in adulthood remain few, excepted in humans for whom effects of parental age are variable according to statistical models or socioeconomic environments. In a small primate which the longevity reaches up to 12 years, we investigated the effects of parental age at conception on the longevity of offspring (N = 278) issued from parents with known longevity. None of the postnatal parameters, including body mass, influenced offspring’s longevity. Mothers’ age at conception significantly affected offspring’s longevity in males but not females. By contrast, fathers’ age at conception did not influence offspring’s longevity. When considering the age of the breeding pairs at conception, the older the mother, the lower the longevity of male offspring with a minimum when the father was aged. No such correlation was observed for female offspring. Lastly, the longevity of female offspring was significantly positively related to the longevity of both parents. Compared with current studies, the surprisingly minor effect of fathers ‘age and the major impact of mothers’ age on male offspring survival, despite favourable conditions, were discussed in relation with the highly seasonal reproduction in mouse lemurs.

## Introduction

Numerous studies have shown a negative effect of advanced parental age on the longevity of offspring. The longevity of offspring of older parents is considered to be shorter than that of the offspring of younger parents. This phenomenom referred as the “Lansing effect” has been decribed in many taxa including more than 300 species [1,2]. However, the majority of results confirming the “Lansing effect” refer to juveniles’ survival, which is mainly explained by an inadequate parental care of aged parents. Depending on the father’s role in parental care, sex differences exist for the effects of mothers ‘age on juveniles’ survival. In birds, juveniles’ survival may be favored by older females, but it also largely depends on the resources provided by the father [3–7]. In fact, the influence of parent’s age can be reduced, even absent, when the environmental conditions are favorable. In mammals, most of studies refer to the negative effects of mothers’ age on the pre-adult survival [2], whereas a few studies reported the role of the paternal age at conception on offspring survival [8, 9].

In humans, numerous recent studies have tested the “Lansing effect” on pre-industrial populations [10, 11] or on modern societies [12–14]. Despite extremely large samples (from to more than five millions of people), contradictory results on the effects of parental age on the longevity of children were described, either negative or positive, depending on the statistical models, on the methods used or on the socio-economic environment to which the lengthening human longevity plays a preponderant role.

Among non-human primates, it is generally considered that pre-adult survival decreases as the mother ages. Among Malagasy species, juvenile survival until weaning is maximal when mothers are middle-aged but rapidly decreases reaching less than 20% for oldest females [15–17]. However, few studies questioned the relationship between parent’s age and longevity of offspring at adulthood.

To test the “Lansing effect” we focused on the grey mouse lemur, a Malagasy primate. In captivity, its longevity may reach up to 12 years [18] with a median lifespan averaging 5.5 years [19]. Mouse lemurs are strict long-day photoperiodic breeders [20]. At the beginning of the breeding season, females enter oestrus and males compete to priority access to females [21]. The male that sired the offspring is therefore determined by both behavioural observations and genetic analyses of paternity. Females give birth to 1 to 3 offspring after a 2-month gestation period and nurse infants for approximately 40 days without male parental care. Records of ages at conception for both mothers and fathers and the relative long longevity of captive mouse lemurs give the opportunity to test the “Lansing effect”.

Using a large database on captive mouse lemur’s life history traits, the aim of this study was to examine whether parental age at conception interfere with the longevity of offspring and potentially whether there exists a relationship may exist between parents’ and offspring’s longevity.

## Material and Methods

### Animals

To investigate the effects of parental age at conception on offspring lifespan, we analysed the longevity data of mouse lemurs for which the age of both parents at the time of reproduction was known. Data were recorded in the mouse lemur history traits database from a laboratory breeding colony established in Brunoy (UMR 7179 MNHN-CNRS, IBISA Platform, agreement F91.114.1, DDPP Essonne, France) from a stock originally caught near the southern western coast of Madagascar sixty years ago.

Captive conditions were maintained constant with respect to ambient temperature (24-26°C) and hygrometry (55-60%). Animals were fed *ad libitum* on a standardized diet, including fresh fruits, a homemade milky mixture (19.3% proteins, 17.2% lipids and 63.5% carbohydrates) and mealworms. To ensure seasonal reproductive rhythms, animals were routinely exposed to an artificial photoperiodic cycle consisting of 6 months of summer-like photoperiod (LD = 14 h of light/day) followed by 6 months of winter-like photoperiod (SD = 10 h of light/day) [20]. The beginning of the breeding season was induced by the exposure to long days. At the time of LD exposure, groups of 2-3 unrelated males with 1 to 3 females were randomly constituted. Immediately, males entered competition for priority of access to oestrous females, leading to a hierarchy mostly depending on aggressive interactions [22].

During sexual competition, several nest boxes were provided so that animals can escape agonistic interactions from conspecifics and the chase or fight immediately stops when the chased animal enters a nest-box. In the colony, heterosexual groups were restricted to the few days of female estruses. Paternity determinations, recorded in the database, allowed calculating the reproductive success of each male within a group. After mating, males were kept in single-sex groups, and females were isolated for gestation, birth and lactation. Litter size and composition were recorded at birth.

### Data analysis

Our analysis was focused on parents (132 mothers and 122 known fathers) and their offspring (140 males and 138 females) that died naturally.

Paternities have been previously determined by genetic analyses and were registered in the database. Briefly, DNA samples were collected from ear or skin tissue samples, extracted (with a QIAmp DNA Mini Kit no. 51306 - Qiagen) and amplified. Genetic analyses were then conducted using random amplified polymorphic DNA method or microsatellite loci [22–24].

To assess which parameters may influence the longevity of offspring, several parameters for both parents and offspring were selected.

First, age at conception for both parents was considered. Mothers’ age at conception averaged = 2.7 ± 0.1 years (N = 278), the majority of mothers being less than 5 years old; mothers aged more than 5 years old represented only 8%. Several females have had several litters with different sires (N = 19). Fathers’ age at conception was significantly higher than that of mothers (μ = 3.2 ± 0.1 years, df_1/554_, F= 16.0, P< 0.001) and 85% of fathers were less than 5 years old. Parents could have offspring from the first breeding season (minimum 260 days), and reproduce until death (no menopause or andropause in mouse lemurs). In our sample however, due to management reasons, parental age at breeding did not exceed 8 years. In this study, parents were considered young when ≤ 2 years old, adults when > 2 to 5 years old, and aged when > 5 years old. Age of the breeding pair corresponded to the sum of the age of both parents, leading to young pairs ≤:4 years (N = 84), adult pairs >4 to 10 years (N=175) and old pairs > 10 years (N=19). Second, parents’ lifespan was included in the analyses.

For offspring, mothers’ parity was not included because previous studies demonstrated that this parameter did not influence the size or the composition of the litters, as well as growth rate during the first month of life [25]. However, size (from 1 to 3) and composition of litter in which offspring were born: male litters (M, MM, MMM), female litters (F, FF FFF) and mixed-sex litters (MF, MMF, MFF) were considered.

Body mass (g) at 30 days, i.e. near weaning time (N = 267), thus providing a clue on maternal investment, as well as at 2 months after birth (N = 274) were included. Lastly, opportunities for offspring to mate and age and timing (relative to the photoperiodic regimen) of natural death were incorporated in the analyses.

### Statistics

Data are presented as means ± SEM. Statistical analyses included a multi-way analysis of variance using or not a covariate and G tests to test distributions. Multiple pairwise comparisons were made using Tukey’s post hoc test. Relationships between the different factors were tested using linear regression analyses.

### Ethics Statement

All the results in this study did not correspond to experimental procedures but are issued from the exploitation of lifespan data collected in the long-term mouse lemur’s captive population maintained for scientific purposes. We adhered to the Guidelines for the Treatment of Animals in Behavioural Research and Teaching [26] and the legal requirements of the country (France) in which the work was conducted. The colony is established under the authorisation of the Direction Départementale de Protection des Populations (DDPP/022-F91-114-1). All procedures to breed mouse lemurs are conducted in accordance with the European Communities Council Directive (86/609/EEC) and are authorized by the Departmental Veterinary Services (Directive 2010/63/UE - Capacity certificate Préfecture de l’Essonne, 04/03/1995). Specifically, for these arboreal primates, housing conditions include cages equipped with branches, various supports, devices to stimulate foraging, and many nesting boxes allowing the animals to express their entire behavioural repertoire. The health and the well-being of captive animals are regularly checked by the animal care keepers and the veterinarian. To improve animal welfare and the enrichment of breeding conditions, meetings of the welfare unit are held every month. When an animal shows signs of poor conditions or signs of social distress, it is isolated and monitored until fully recovered. Lastly, only animals that were found dead, were used to research on longevity, so no anaesthesia, euthanasia or animal sacrifice were part of this study.

## Results

### 1) Offspring longevity and postnatal parameters

The median lifespan of offspring used in this study reached 5.5 ± 0.1 years (N = 278) without significant difference between males and females (df_1/276_, F=0.12, P = 0.7). Maximal lifespan reached 12.1 years in males and 11.1 years in females. Natural deaths mainly occurred at the photoperiodic transitions (43%, Gdf_2_ = 13.6, P < 0.01) but were significantly more frequent during long-day photoperiod (df_1/274_, F = 4.965, P = 0.03) independently of sex (df_1/274_, F = 0.01, P = 0.9).

#### Size and composition of the litter

Offspring mainly issued from litters of 3 (55%) in mixed sex litters (63%), litters of 1 or 2 babies reaching only 9% and 36% respectively. The distribution of litter size was identical within sexes (Gdf2 =2.13 NS).

Mothers’ age at conception had an impact on the size of the litter produced (r = 0.159, P = 0.008), with young females producing less numerous triplets than older females (45% versus 66% respectively - df_2/275_, F= 3.25.18, P = 0.04). The composition of the litter was independent of the mothers’ age (df_2/275_, F= 2.42, P = 0.09). By contrast, whereas fathers’ age had no impact on litter size (r = 0.092, P= 0.1), younger fathers produced less numerous male-biased litters (df_2/275_, F= 4.3, P = 0.01).

Consequently, the size and the composition of the litter were dependent on the age of the breeding pair with adult pairs producing more numerous offspring (63%) in more numerous mixed-sex triplets (size df_2/275_, F= 4.09, P = 0.02 and composition df_2/275_, F= 5.99, P =0.003)

But offspring longevity was independent of the size (df_2/275_, F= 0.73, P = 0.5) and of the composition of the litter (df_2/274_, F = 0.08, P = 0.7), whatever the offspring sex (df_2/274_, F = 0.08, P = 0.9 - Fig 1).

**Fig. 1 –.**
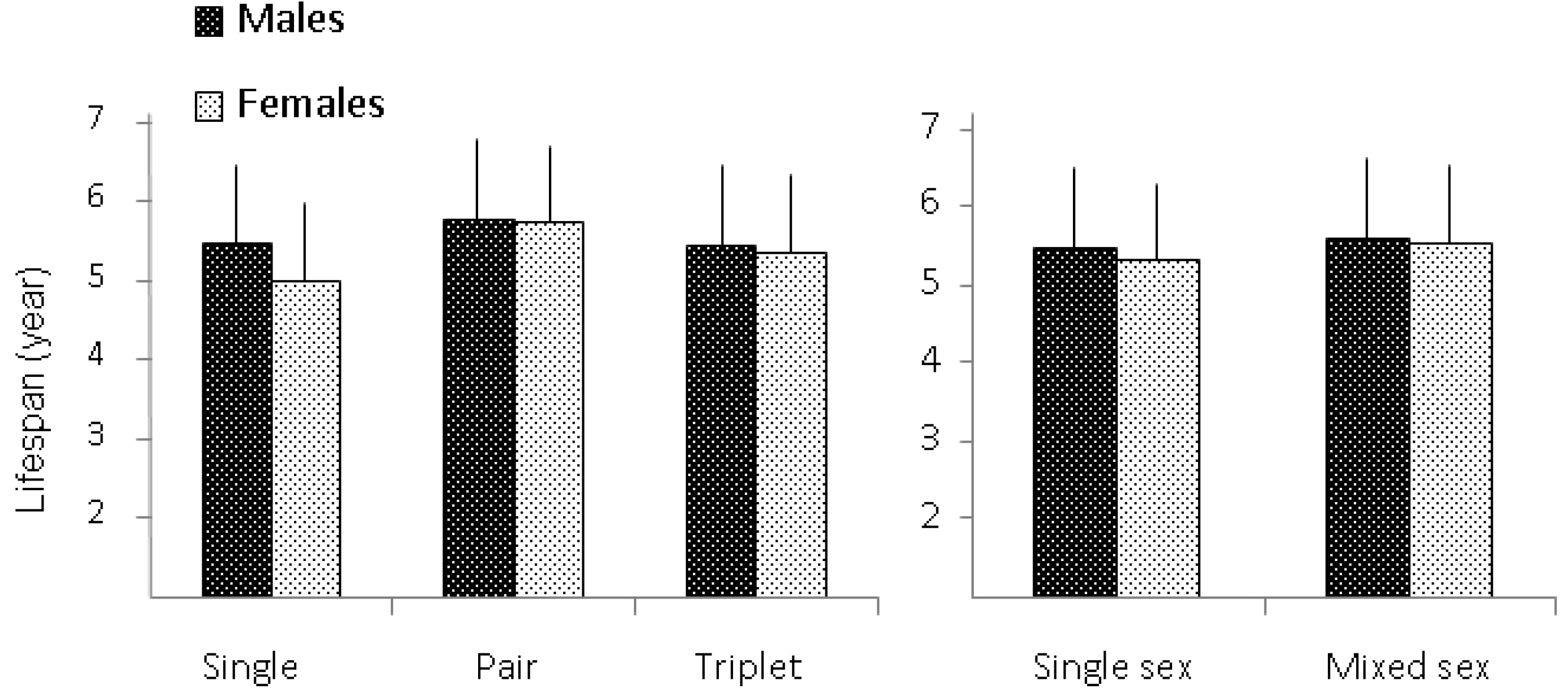
Lifespan (means ± SEM in years) in female and male offspring according to the size (left panel) and the composition (right panel) of the litters.

**Fig. 1 - *Lifespan (means ± SEM in years) in female and male offspring according to the size (left panel) and the composition (right panel) of the litters.***

#### Post natal body mass

Offspring body mass reached at weaning time (30 days) averaged 32.1 ± 0.4g (N = 266 - Table I). It was independent of both mothers ‘age (r = 0.043, NS) and the fathers’ age (r = 0.02, NS), but it was strictly linked to the size of the litter. Indeed, body mass of animals born in triplet was significantly lower (*df*_2/260_, F = 14.1, P < 0.001) than that of animal born alone or in pairs, independently of sex (*df*_1/260_, F = 0.09, P = 0.8) and of litter the composition (*df*_2/263_, F = 1.3, P = 0.2).

**Table I.** - Body mass (g, mean ± E) of female and male offspring at weaning time (30 days) and 60 days after birth depending on litter size. * Significant differences between triplets and other litters (P < 0.001).

| Litter size | Body mass (30 days) |  | Body mass (60 days) |  |
| --- | --- | --- | --- | --- |
|  | Females | Males | Females | Males |
| Single | $31.9 \pm 2.8$ (9) | $34.9 \pm 2.2$ (13) | $50.8 \pm 2.4$ (9) | $58.6 \pm 2.2$ (13) |
| Pair | $35.1 \pm 0.8$ (51) | $33.5 \pm 0.9$ (47) | $53.1 \pm 1.2$ (51) | $52.3 \pm 1.2$ (49) |
| Triplet | $30.4 \pm 0.8$ (75)* | $30.3 \pm 0.7$ (71)* | $49.3 \pm 1.0$ (77)* | $47.9 \pm 0.9$ (74)* |
| Mean | $32.3 \pm 0.6$ (135) | $31.9 \pm 0.6$ (131) | $50.8 \pm 0.8$ (137) | $50.5 \pm 0.7$ (136) |

Offspring body mass averaged 50.6 ± 0.5 (N = 267 Table I) 2 months after birth without effect of sex (P = 0.92). It remained however dependent on the litter (*df*_2/267_, F = 11.6, P < 0.001) but not of litter composition (P = 0.8) although male offspring born without littermates were significantly heavier (df_2/267_, F = 3.31, P = 0.04). Lastly, maternal investment in terms of offspring’s body mass did not depend on mothers’ age at conception (respectively P = 07 at 30 days, P = 0.8 at 60 days)

Offspring longevity was not linked to their body mass at weaning for both males (r = 0.109, P = 0.2 N = 131) and females (r = 0.111, P = 0.2, N = 135). Likewise the body mass reached by offspring 2 months after birth was unrelated to their lifespan whatever the sex (r = 0.078, P = 0.2, N = 267).

### 2) Offspring’s longevity and reproductive investment

Within female offspring, no difference in longevity was observed between females that had access to reproduction at least once and those that did not come in contact with males (respectively 5.12 ± 0.2 years, N = 55 versus 5.62 ± 0.2 years, N = 83 - df_1/136_, F=2.7, P = 0.1). By contrast, males having at least one opportunity to mate had significantly higher longevity than the others (respectively 6.04 ± 0.2 years N= 30 versus 4.26 ± 0.3 years, N= 102, df_1/138_, F=17.6, P <0.001).

### 3) Offspring ‘s longevity and parental age at conception

Mothers’ age at conception was negatively correlated to offspring’longevity for males (r = - 0.309, P < 0.001) but not for females (r = 0.104, P = 0.2 - Fig.2). However, in both sexes, the highest offspring longevities were oberved for mothers younger than 3 years old: 5.9 ± 0.2 years (N =182) versus 4.9 ± 0.2 years for older females (N = 96).

**Fig. 2 –.**
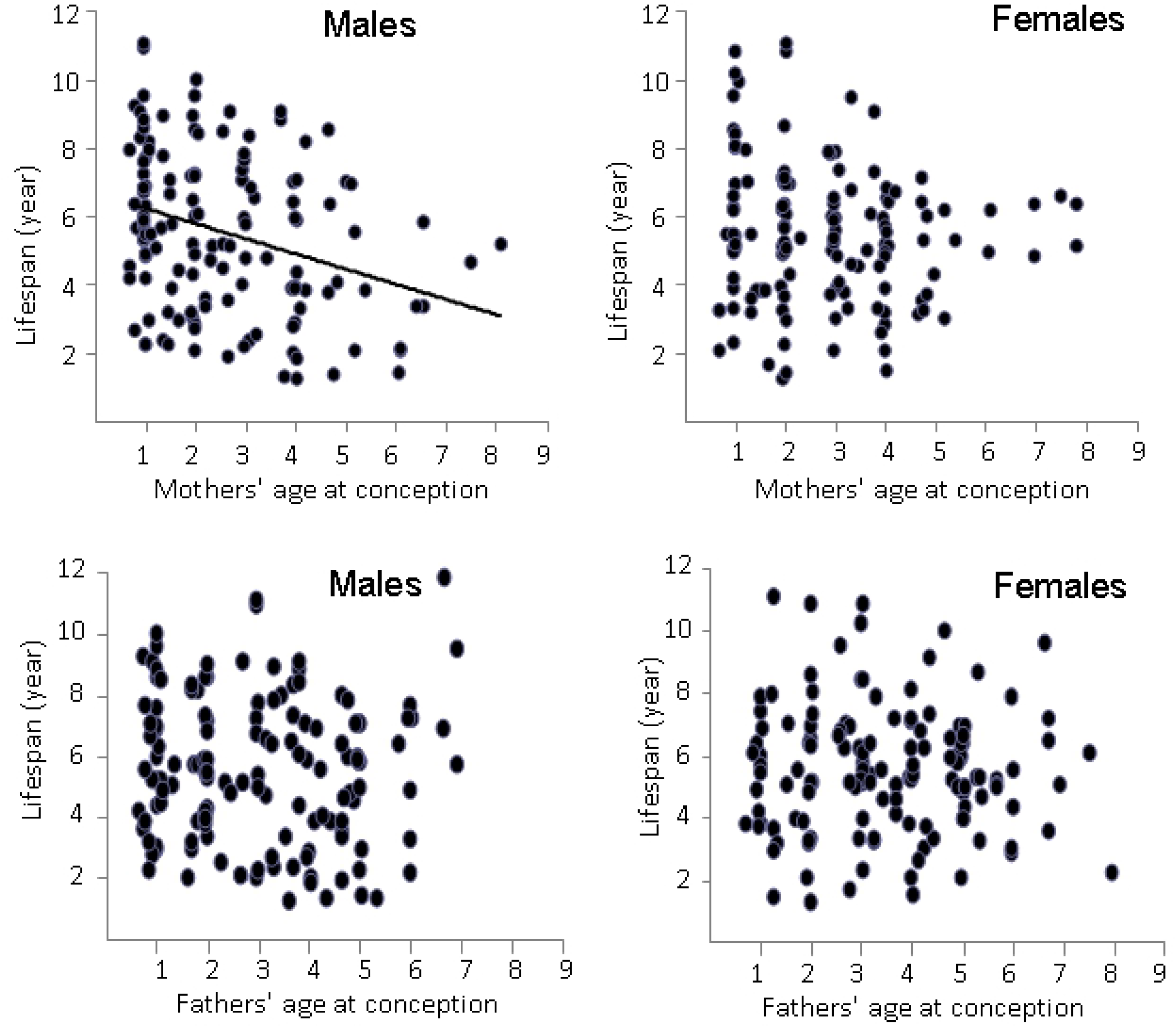
Relationship between offspring longevity in males (left panel) and females (right panel) according to the age at conception of mothers and fathers. A significant correlation was observed between age at conception of mothers and longevity in offspring males (r = - 0.309, P < 0.001).

By contrast, the fathers’ age at conception did not influence offspring’ longevity for both males (r = 0.057, P = 0.5) and females (r = 0.027, P = 0.8 - Fig.2).

**Fig.2 - Relationship between offspring longevity in males (left panel) and females (right panel) according to the age at conception of mothers and fathers. A significant correlation was observed between age at conception of mothers and longevity in offspring males (r = - 0.309, P < 0.001).**

Since fathers’ age at conception did not influence the lifespan of offspring, the effect of the breeding pairs’ composition on offspring’s longevity was mainly dependent of the mothers’ age (Fig.3). The composition of the breeding pair was negatively correlated to the lifespan of offspring males (r = - 0.240, P = 0.004) but not of offspring females (r =-0.082, P = 0.3). As more as the breeding pair aged, the longevity of male offspring decreased (r = 0.2400, P < 0.01) with a minimum lifespan when fathers were aged (P< 0.02). In addition, when analysing the difference in parental age at conception, the longevity of male offspring was significantly shorter when mothers were older than fathers (r = 0.1884, N=140, P <0.05). No such correlations were observed for female offspring (mother r = 0.06, and father r = 0.08. NS).

**Fig. 3 –.**
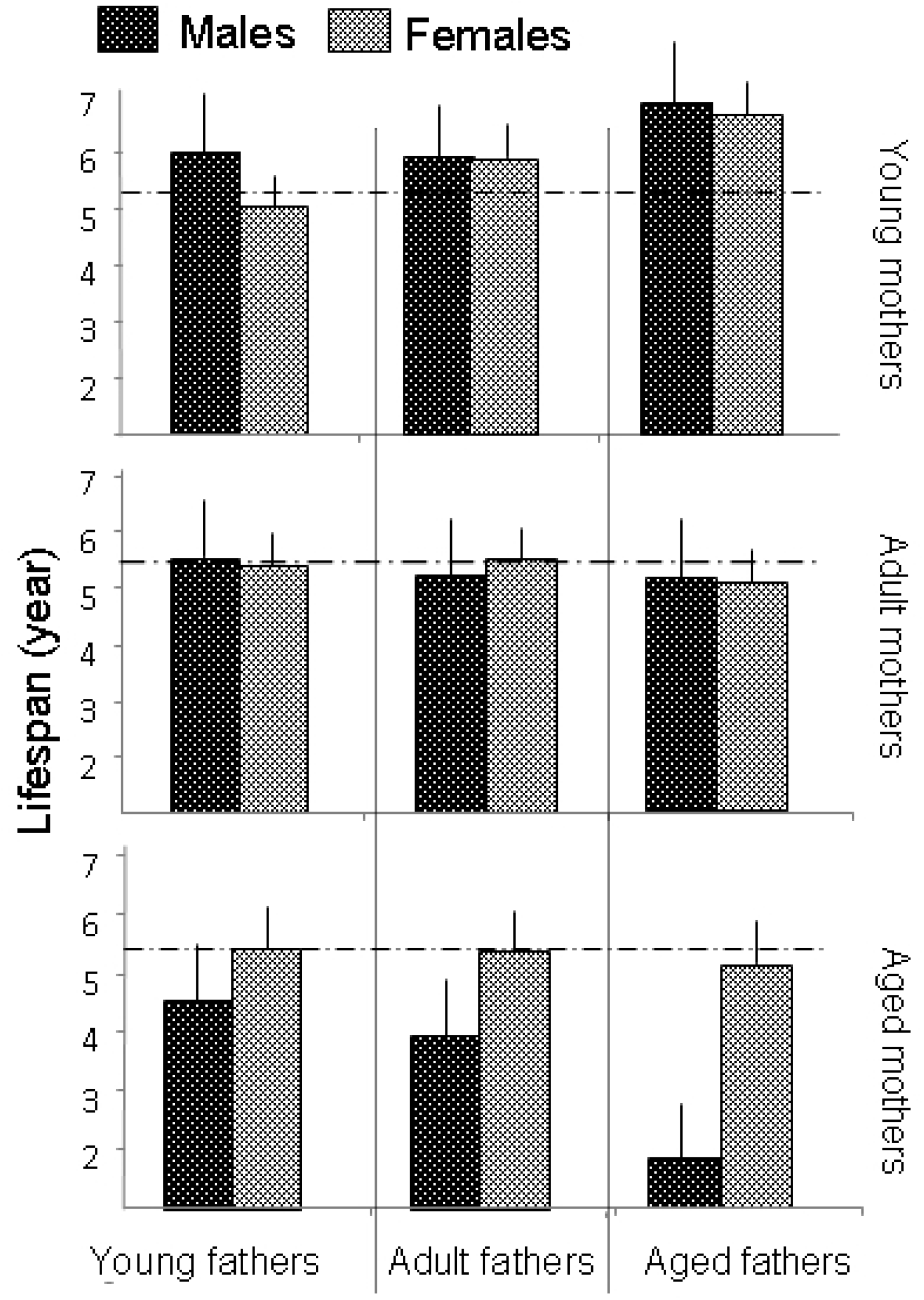
Lifespan (means ± SEM) of male and female offspring according to the composition of breeding pairs. The broken line indicates the mean survival (5.5 years).

**Fig.3 - Lifespan (means ± SEM) of male and female offspring according to the composition of breeding pairs. The broken line indicates the mean survival (5.5 years).**

Lastly, when considering the potential relationship between parental longevity and that of their offspring, a significant difference appeared according to sex. The lifespan of males offspring was unrelated to the longevity of their respective mother (r = 0.006, P = 0.9) and father (r = 0.078, P = 0.3). By contrast, the longevity of offspring females correlated significantly with both their mother (r = 0.1682, P = 0.04) and their father (r = 0.2133, P = 0.01).

## Discussion

The longevity of captive mouse lemurs may reach up to 12 years with a 50% survival time around 5.5 years [18, 19] and with no significant difference between sexes. Mouse lemurs are highly seasonal breeders and can reproduce throughout all their life even at an advanced age. Results demonstrated that mothers’age at conception affected only the longevity of males offspring while the fathers’ age at conception had no influence on longevity of offspring, whatever their sexes. As more as the mother aged, the survival of male offspring’s decreased with a minimum when the father was aged. More, influence of parents longevity in females was already described in humans [10, 27, 28], which is not the case for .male offspring.

Among vertebrate species, the age of mothers for the pre-adult survival has been well documented [1, 2]. With ageing, a decrease in fertility and in maternal investment is considered to be the key for reduced survival of offspring. In the very good conditions of captiviity, age does not affect either fertility or maternal investment in female mouse lemurs. As mother ages, litter size increases and postnatal growth of offspring remains unaffected. Thus, ofspring survival in adulhood is not dependent of an inadequate maternal investment or on postnatal conditions in captive mouse lemurs.

In our sample, fathers’ age at conception had no impact on offspring longevity whatever the sexes. Thus, the maternal contribution to the offspring survival may out-weight the paternal component and affect mostly male offspring longevity. This strongly suggests a predominant role of genetic load from the mother.

The effect of parental age on offspring longevity is generally attributed to a direct age-related deterioration of the germ cells: DNA mutation, DNA methylation, shortening of telomeres [29]. Within age-related deteriorations, the telomere length (TL) is recognized as the most suitable biomarker of aging [30] and telomere inheritance appears to be paternal in mammals [31]. A large comparative study on mammalian species, including humans, showed clear relationships between age-related shortening of TL and lifespan [31, 32]. However, the age-related TL shortenning appears to be sex- and species-dependent [33–36]. If strong evidence exists for genetic inheritance of TL, conclusions on relationship between parental age at conception and TL of offspring depend on the species studied. In several studies (mostly birds), TL shortens as parents age and predicts offspring TL with sex-specific differences in TL inheritance [4, 6, 37–41]. By constrast, a weak or no relationships between TL and parental TL has been observed in other species [42–45].

Telomere biology in mouse lemurs seems an exception among primates. The average telomere length is among the longest reported for primate species and, no detectable TL shortening with ageing was detectable [46]. Likely to hamsters during short day period [47] mouse lemurs use daily torpor that may contribute to an increase in TL. More, in small-bodied species, the seasonal variation in sperm production might explain the lack of an effect of parental age on TL [43]. In male mouse lemurs, a greater sperm production, the presence of telomerase activity in testes and social dominance required for successful mating may be another factor for maintenance of TL [48, 49]. Dominant males (i.e. fathers) would invest more heavily in soma reparation and TL maintenance as suggested by the longer lifespan of males that were offer the opportunity to reproduce at least once. All these characteristics could explain the lack of effect of fathers’ age at conception on offspring longevity. Similarly, for female mouse lemurs, the use of torpor, the high reproductive seasonality and the protective role of oestrogens could contribute to TL maintenance over life.

Why mothers’ age at conception affects only male offspring remains however a question. As supported by Entringer et al. 2015 [50], mothers’ oestrogens levels of the mother would predict TL of offspring. With ageing, levels of oestrogens in female mouse lemurs significantly decrease and male-biased litters are more frequent [51]. But, this decrease does not impact females’ fecundity or female offspring longevity. TL, mitochondrial DNA content is higher in females and no deterioration was observed at advanced maternal age [52]. Thus, male offspring from aged mother would not suffer from lesser portion of maternal genetic load. In our study, the lifespan of male offspring was reduced when the mother was older than the father. This suggests interplay between the TL of the father and the age-related deterioration of maternal genetic load if any. However, the sex-specific effect of aged mothers on offspring longevity could rely to the relative low number of male offspring issued from aged breeding pairs. More parameters would deserve to be further explored such as the composition of the litters [53], age-related maternal behaviours or social interactions later in the weaning period.

According to several studies [10, 54, 55], life-history trajectory of offspring could vary according to mothers’ age suggesting potential trans-generational effects of maternal age. For female mouse lemurs, it has been already described that, in the highly favourable conditions of captivity, there was no cost of reproduction for longevity [56]. In several species, offspring from older mothers show changes in their lifetime reproductive success, associated or not with a shorter lifespan [54, 57, 58]. Further work is needed to know whether reproductive success in female mouse lemurs is affected by maternal age at conception. It would be of interest to follow the longevity of all female offspring from a maternal line to decipher the inheritance of female longevity.

## Acknowledgements

We are extremely indebted to I. Hardy for creating the “*Mouse lemur life history traits*” database (1995 - present) and to Dr J. Terrien for English editing. We would like to thank L. Dezaire, S. Gondor and I. Hiron, the animal caretakers, for their contribution to the excellent care of the mouse lemurs.

